# Alterations in striato-thalamo-pallidal intrinsic functional connectivity as a prodrome of Parkinson’s disease

**DOI:** 10.1101/140830

**Authors:** Eran Dayan, Nina Browner

## Abstract

Although the diagnosis of Parkinson’s disease (PD) remains anchored around the cardinal motor symptoms of bradykinesia, rest tremor, rigidity and postural instability, it is becoming increasingly clear that the clinical phase of the disease is preceded by a long period of neurodegeneration, which is not readily evident in terms of motor dysfunction. The neurobiological mechanisms that underpin this prodromal phase of PD remain poorly understood. Based on converging evidence of basal ganglia (BG) dysfunction in early PD, we set out to establish whether the prodromal phase of the disease is characterized by alterations in functional communication within the input and output structures of the BG. We analyzed resting-state functional MRI data collected from patients with REM sleep behavior disorder (RBD) and/or hyposmia, two of the strongest markers of prodromal PD, in comparison to age-matched controls. Relative to controls, subjects in the prodromal group showed reduced intra- and interhemispheric functional connectivity in a striato-thalamo-pallidal network. Functional connectivity alterations were restricted to the BG and did not extend to functional connections with the cortex. The data suggest that local interactions between input and output BG structures may be disrupted already in the prodromal phase of PD.

## Introduction

While advances in our understanding of Parkinson’s disease (PD) keep accumulating, its diagnosis remains centered around the presentence of cardinal motor symptoms, including bradykinesia, rest tremor, rigidity and postural instability (Mahlknecht et al., 2015; Postuma and Berg, 2016). Nevertheless, it is by now clear that these symptoms are preceded by a long period in which neurodegeneration spreads throughout the nervous system (Postuma and Berg, 2016). Indeed, multiple lines of evidence point to the existence of a prodromal phase in PD, where various symptoms are present, but are not yet sufficient to define the disease or to meet established diagnostic criteria (Mahlknecht et al., 2015; Postuma and Berg, 2016). Markers for the prodromal phase of PD are still under investigation. Studies have suggested that a polysomnography-based diagnosis of rapid eye movement (REM) sleep behavior disorder (RBD) is among the strongest markers (Berg et al., 2015; Mahlknecht et al., 2015; Postuma et al., 2015a), since up to 80% of patients with RBD will eventually develop a neurodegenerative disease, primarily PD (Postuma and Berg, 2016). Other markers commonly cited in the literature include hyposmia, and a range of autonomic dysfunctions (Mahlknecht et al., 2015; Postuma et al., 2015a).

Little is known to date about the neurobiological mechanisms that characterize the prodromal phase of PD. An influential staging scheme for PD postulated that α-synuclein pathology first begins in the caudal brainstem and the olfactory bulb, later progressing to the substantia nigra (SN) and finally to limbic regions and the neocortex (Braak et al., 2003). However, it remains unclear if any of these regions and pathways are affected during the prodromal phase of PD. Neuroimaging provides a useful non-invasive means of addressing this question. In particular, altered functional connectivity between the SN and the putamen and SN and parieto-occipital cortical regions were reported in patients with RBD, when compared to controls (Ellmore et al., 2013). More recently reduced resting-state coactivation within a large-scale network of brain regions including the putamen, middle, medial and orbital frontal cortices was found in patients with RBD compared to age-matched controls (Rolinski et al., 2016). Together, these two studies suggest that functional interactions between the putamen and other cortical and subcortical regions are disrupted in the prodromal phase of PD. However, findings from other imaging modalities suggest that other regions in basal ganglia (BG) may be affected (Postuma and Berg, 2016). For example, molecular imaging studies have found alterations in dopaminergic neurotransmission in patients with idiopathic RBD in both the putamen and the caudate (Eisensehr et al., 2000; Iranzo et al., 2010). These findings join data on aberrant functional connectivity (Rolinski et al., 2015) and regional atrophy (Zeighami et al., 2015) in early PD found in the caudate and pallidum, as well as in the putamen.

In the classical model of BG organization (for a recent review, see Lanciego et al., 2012), cortical signals flow through the striatum, forming two pathways. In the direct pathway, output from the striatum reaches directly to the substantia nigra pars reticulata (SNr) /internal globus pallidus (GPi). In the indirect pathway, striatal output reaches the SNr/GPi via the external globus pallidus (GPe) and the Subthalamic nucleus (STN). Signals from the SNr/GPi then project back to the cortex through the thalamus. Complex inhibitory and excitatory interactions underlie the normal functioning of this intricate circuitry, and alter in movement disorders such as PD (Lanciego et al., 2012). Still, the time course of neuropathological changes within the BG, and in particular the extent to which intrinsic functional connectivity between the different structures in this circuity is altered in the prodromal phase of PD remains insufficiently understood. Here, we set out to examine if the prodromal phase of PD is associated with specific alterations in functional connectivity within the basal ganglia and between the BG and other cortical and subcortical regions.

## Materials and Methods

### Subjects

Data from 35 participants were analyzed, including 17 participants in a prodromal PD group (mean age= 67.88 ± 4.8, 12 males) and 18 age-matched controls (mean age 64.22 ± 9.77, 14 males). The data were obtained from the Parkinson’s Progression Markers Initiative (http://www.ppmi-info.org), a comprehensive observational, international, multi-center study designed to identify PD progression biomarkers (Marek et al., 2011). All participants provided written informed consent and the procedures were approved by the Institutional Review Boards of the participating centers. All resting-state and anatomical scans provided by the PPMI for both groups were included in the analysis. To be included in the prodromal PD group participants had to be at least 60 years of age and present at least one of the two following clinical characteristics: a) hyposmia, as confirmed using the University of Pennsylvania Smell Identification Test (UPSIT)(Doty et al., 1984), with a cutoff at or below the 10th percentile adjusted by age and gender. b) polysomnography meeting the criteria of RBD and/or a clinical diagnosis of RBD by site investigator including existing polysomnography (PSG). Current or active clinically significant neurological or psychiatric disorders including a current diagnosis of PD were defined as exclusion criteria in this group. A total of 4 subjects in the prodromal group met the criteria for hyposmia and 13 met the criteria for RBD. For the control group, current or active clinically significant neurological disorder or a first degree relative with idiopathic PD were defined as exclusion criteria. Full inclusion and exclusion criteria for the two groups are available online (http://www.ppmi-info.org). Data from 2 subjects in the prodromal group (both were RBD patients) and 1 in the control group were excluded because of excessive head motion during scanning (see bellow), therefore the final dataset included 15 participants in a prodromal group (mean age=68.06±5.07, 11 males) and 17 age-matched controls (mean age=63.64 ±9.75, 14 males).

### Cognitive and motor evaluation

As part of the PPMI protocol, subjects underwent basic screening for cognitive function, using the Montreal Cognitive Assessment (MOCA, score range: 0-30) (Nasreddine et al., 2005). Subjects also underwent a detailed exam using the unified Parkinson’s disease rating scale (UPDRS). Scores from part III of the UPDRS, focusing on motor dysfunction (range: 0 to 108) were extracted for each subject as a measure of motor function.

**Table 1.**
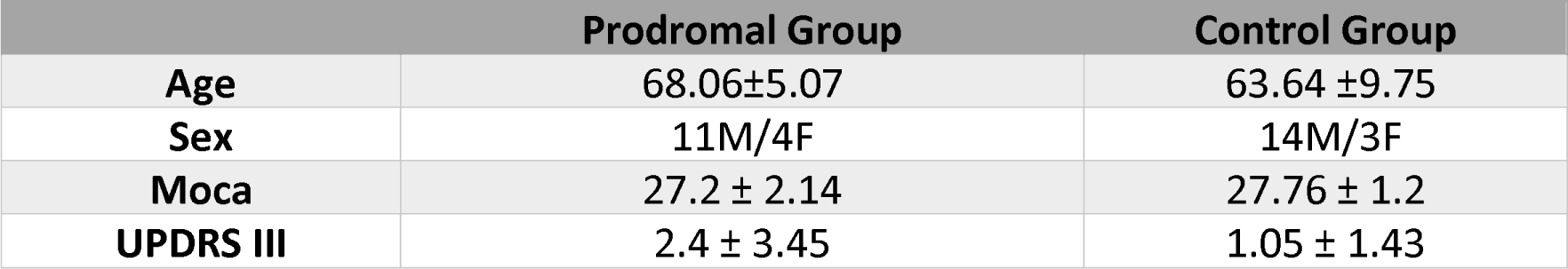
Demographic and clinical characteristics in the prodromal and control groups.

|  | Prodromal Group | Control Group |
| --- | --- | --- |
| **Age** | 68.06±5.07 | 63.64 ±9.75 |
| **Sex** | 11M/4F | 14M/3 F |
| **Moca** | 27.2 ± 2.14 | 27.76 ± 1.2 |
| **UPDRS III** | 2.4 ± 3.45 | 1.05 ± 1.43 |

### Imaging data acquisition

Imaging data were acquired with Siemens 3T scanners (either Prisma or TimTrio). Structural images were acquired with an MPRAGE sequence using GRAPPA (TE=2.98, TR=2300 ms, FA=9°, 1 mm^3^ isotropic voxel). Resting-state functional MRI (fMRI) scans were acquired from the same subjects with an echo-planar sequence (total of 210 volumes [164 in 2 of the prodromal subjects), 40 axial slices, acquired in ascending order, TE=25.0 ms; TR=2400.0 ms, Flip Angle=80.0° voxel size = 3.3 mm3 [3.0 mm in 2 of the prodromal subjects]. For the resting state scans subjects were instructed to rest quietly with their eyes open while trying not to fall asleep.

### Imaging data analysis

Data analysis was performed using SPM12 (http://www.fil.ion.ucl.ac.uk/spm/software/spm12/) and the Conn toolbox (Whitfield-Gabrieli and Nieto-Castanon, 2012) version 16b, both running on MATLAB R2016b (MathWorks, Natick, MA, USA). Functional images were realigned, unwarped and slice-time corrected; grey-matter, white-matter and cerebrospinal fluid (CSF) were segmented and the functional data were normalized to the MNI template. Motion artifacts were further detected using the ART-based scrubbing method (see, Power et al., 2012), implemented in Conn (setting a threshold of 2 mm subject motion and a global signal threshold of Z=9) and the data were spatially smoothed with a Gaussian kernel set at 8mm full width at half maximum. Unsmoothed data was used in the region-to-region analysis (see bellow). Signals from white matter and CSF (the first five principal components from each), as well as the 6 motion realignment parameters and their first order derivatives were regressed out of the signal, and so were outlier volumes (including the single volumes preceding the outliers) detected in the ART-based scrubbing procedure. The data was also linearly detrended. Data from 2 subjects in the prodromal group and 1 subject in the control groups were excluded, because of excessive head motion (>50% of volumes detected as outliers), leaving 15 subjects in the prodromal groups and 17 in the control group. The residual signals were then band-pass filtered (0.008Hz to 0.09Hz).

Two types of analyses were conducted. First, interactions between regions of interest (ROIs) in the BG were tested with region-to-region analysis. In the first level of analysis region-to-region connectivity matrices were computed for each subject. Our specific focus in the current study was on connectivity within BG circuitry. We thus defined ROIs in the caudate, putamen and thalamus based the probabilistic Harvard-Oxford atlas. Additional ROIs in the SN, STN, GPi and GPe were defined based on an age-appropriate BG atlas (Keuken et al., 2017). In the ROI-to-ROI analysis, each subject’s fisher-Z transformed connectivity matrices, expressing pairwise correlations between the blood-oxygenation level-dependent (BOLD) time series of each pair of ROIs, were subjected to a second-level analysis testing differences between the prodromal and control groups. The analysis was based on a connection-level threshold set at False Discovery Rate (p-FDR)⍰<⍰0.05, and an intensity-based family-wise error (FWE) correction (p-FWE < 0.05), based on the non-parametric network-based statistics (NBS) approach (Zalesky et al., 2010). Briefly, suprathreshold pairwise connections where differences between the two groups existed were detected and their intensity was computed. Then, random permutations where subjects were randomly divided into 2 groups were performed, retaining the overall strength of suprathrshold connections in each permutation. The overall procedure yields a null distribution of connection strengths which can then be used to identify a subgroup of connections in the original network that survive FWE correction.

Second, functional connectivity between the BG and other cortical and subcortical regions was additionally tested using a seed-to-voxel approach. In the first level of the analysis, seed-to-voxel maps were computed for each subject, with seed regions comprising right and left putamen, both defined based on the Harvard-Oxford atlas. A secondary analysis also examined seed-to-voxel functional connectivity, based on seed ROIs in right and left SN, defined based on a standardized age-appropriate BG atlas (Keuken et al., 2017). The single-subject seed-to-voxel maps were subjected to a random effect second-level general linear model testing for differences between the groups. A voxel-level threshold of p>0.001 was used, p-FDR (>0.05) corrected for multiple comparisons at the cluster level.

## Results

Our objective was to examine if intrinsic functional interactions within the BG and between the BG and other cortical and subcrotical regions differ in patients who are suspected to be in the prodromal phase of PD, relative to age-matched controls. We analyzed resting-state fMRI data acquired from a group of individuals who had a confirmed diagnosis of RBD and/or were hyposmic, two established markers of prodromal PD (Postuma and Berg, 2016). Data acquired from age-matched controls was used as a comparison.

The prodromal and control groups did not differ in age (t_30_=1.574, p>0.1), male/female distributions (Kolmogorov-Smirnov Z=0.255, p>0.1), cognitive function, assessed with the MOCA (t_30_=-0.934, p>0.1) or motor function as assessed with UPDRS part III (t_30_=1.464, p>0.1). In addition, we did not find statistically significant differences between the groups in the total amount of head movement during scanning (t_30_=1.467, p>0.1) or the number of outliers detected in the ART-based scrubbing procedure (Mann-Whitney U=117, p>0.1), ruling out possible confounds which may have been introduced by head motion during scanning.

### Region-to-region functional connectivity in the basal ganglia

To identify possible functional connectivity alterations within the BG found in the prodromal group, relative to the control group we first subjected the bilateral putamen, caudate and pallidum ROIs to network-based statistics analysis (Zalesky et al., 2010). Functional connectivity within the BG was first derived for the prodromal (Fig 1a), the control (Fig 1b) groups. A comparison between the groups (with a connection-level threshold set at p-FDR⍰<⍰0.05 and FEW network-level correction set at p-FWE < 0.05) revealed that, relative to controls, the prodromal group displayed reduced STN-striato-thalamo-pallidal inter- and intrahemispheric functional connectivity (Fig. 1c). The majority of connections differentiating the two groups were in striato-thalamo-pallidal (primarily the GPe) functional connections. Increases in functional connectivity in the prodromal group relative to the control group were not detected under the same thresholds. Connectivity in the BG network did not correlate with the prodromal group’s MOCA (r=-0.1, ns) or UPDRS scores (r=0.13, ns). Altogether, connectivity in the striato-thalamo-pallidal network identified above differentiated the prodromal and control groups with a Sensitivity of 93.33% (95% confidence interval (CI) of 68.05% to 99.83%) and Specificity of 82.35% (CI of 56.57% to 96.20%).

**Figure 1.**
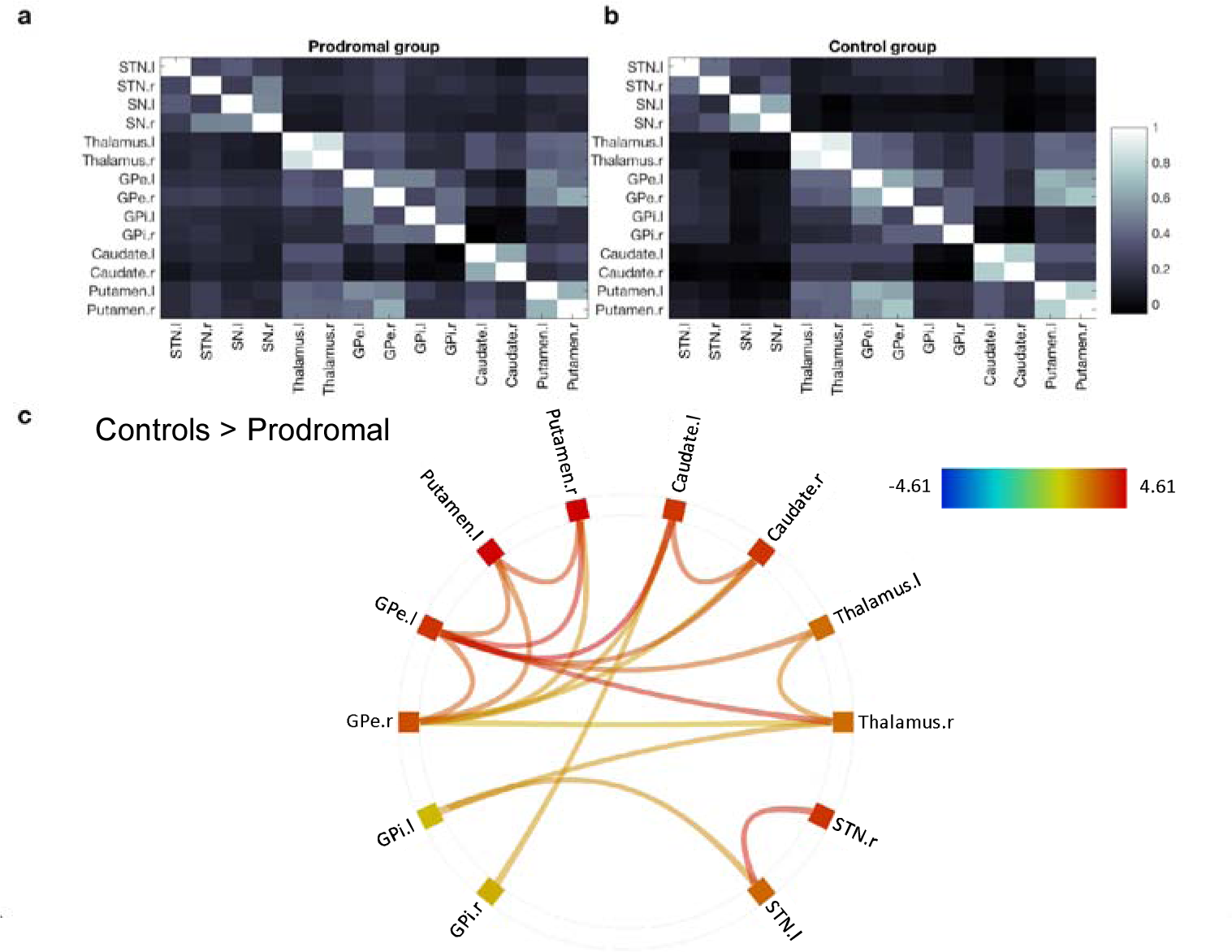
Region-to-region functional interactions in the basal ganglia. (a) Average functional connectivity within the basal ganglia network in the prodromal group (b). Average BG functional connectivity in the control group. All correlation values were inverse Fisher Z transformed (to Pearson’s r values) for visualization purposes. (c) Functional connections for which the prodromal group showed reduced connectivity relative to the control group, based on a non-parametric network-based statistics analysis (family-wise error correction, p-FWE⍰<⍰0.05) coupled with a seed-level threshold set at p-FDR ⍰<⍰0.05. Shown are pairs of network nodes between which the resting-state times series correlations were weaker in the prodromal group than in the control group. Colored lines depict the strength of the test statistic.

### Seed-to-voxel functional connectivity in the basal ganglia

The results above suggest that an extensive set of intrinsic functional connections within the BG differentiated the two groups. We next tested whether the intrinsic alterations in functional connectivity found in the prodromal group, relative to the control group, were restricted to the BG, or rather were also evident in interactions between the BG and other cortical and subcortical regions. We therefore extracted the fMRI signals from right and left putamen and used these as seed regions in a seed-to-voxel analysis. We focused on functional connectivity with the putamen for two reasons. First, the putamen as its interactions with other cortical and subcortical regions have been consistently shown to be disrupted in the prodromal phase of PD (Ellmore et al., 2013; Rolinski et al., 2016). Second, as cortical input to the BG passes through the striatum, disruptions in cortical-BG interactions, if such indeed exist, should be reflected in functional connectivity between cortical regions and the putamen. The results of this analysis suggest that BG connectivity differences between the prodromal and the control groups were primarily limited to functional interactions within the BG. The prodromal group showed reduced functional connectivity between the right putamen seed and a cluster encompassing contralateral putamen and pallidum (p<0.0001, FDR corrected at the cluster level). This functional connection differentiated the two groups with a Sensitivity of 80.0% (CI 51.91% to 95.67%) and Specificity of 82.35% (CI 56.57% to 96.20%). The prodromal group also showed reduced functional connectivity between the left putamen seed and a cluster encompassing bilateral pallidum, putamen and caudate (p< 0.0001, FDR corrected at the cluster level). This functional connection differentiated the two groups with a Sensitivity of 86.67%% (CI 59.54% to 98.34%%) and Specificity of 88.24 % (CI 63.56% to 98.54%). No clusters showed increased functional connectivity in the prodromal group relative to the control group.

**Figure 2.**
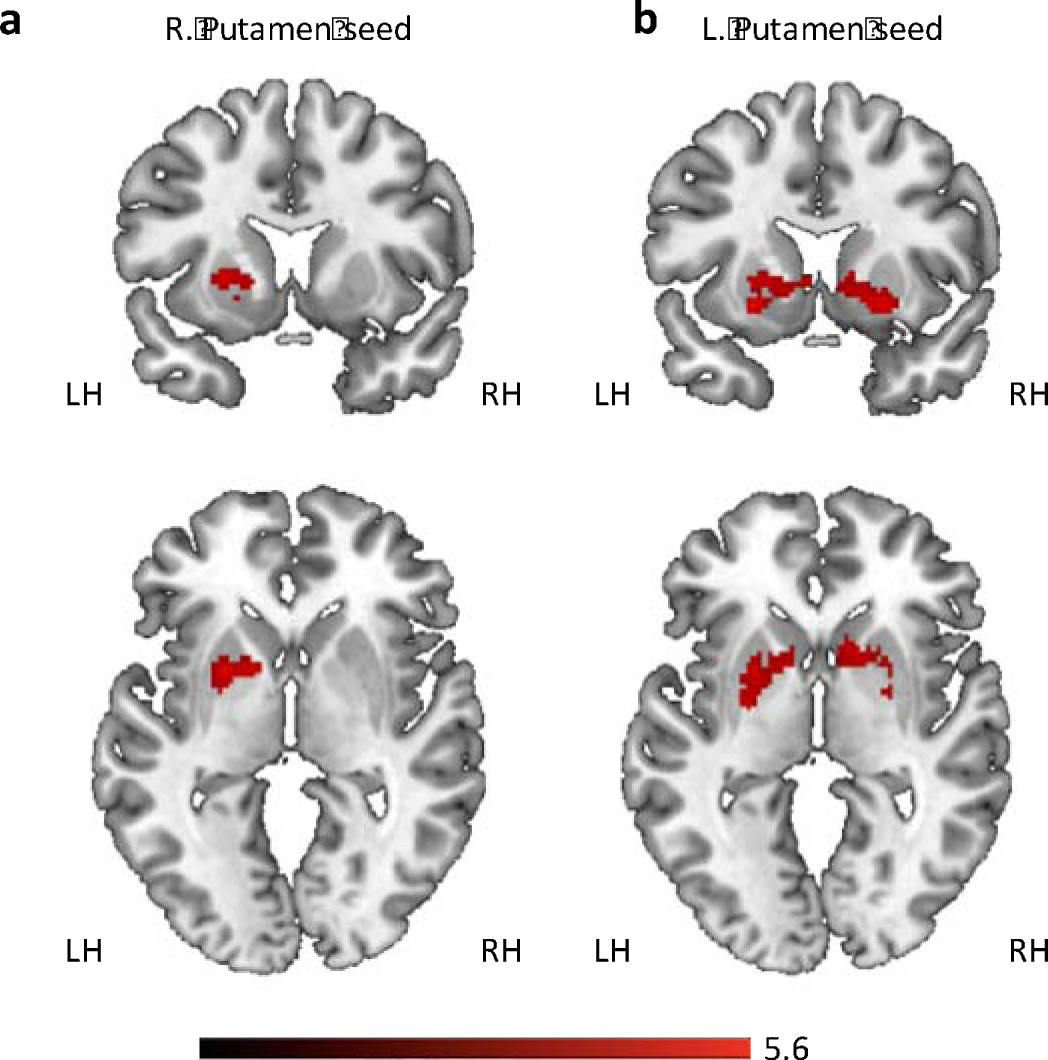
Seed to voxel functional connectivity analysis. (a) Relative to the control group, the prodromal group displayed reduced functional connectivity between right putamen and a cluster encompassing left putamen and pallidum. (b) Reduced functional connectivity in the prodromal relative to the control group was also found between left putamen and clusters in the left pallidum, and right putamen and pallidum. LH, left hemisphere. RH, right hemisphere.

**Table 2.** Seed-to-voxel analysis. Clusters showing reduced functional connective in the prodromal group, relative to the control group.

| Cluster X,Y,Z | Locations | Cluster size | Cluster p-FDR | Peak p |
| --- | --- | --- | --- | --- |
| *R. Putamen seed* |  |  |  |  |
| -22, 6, 2 | Putamen (L), Pallidum (L) | 359 | 0.0012 | <0.0001 |
| *L. Putamen seed* |  |  |  |  |
| 24, 12, -8 | Putamen (L,R), Caudate (L,R) Pallidum (L,R) | 654 | <0.0001 | <0.0001 |

### Functional connectivity with the substantia nigra

Previous results suggested that functional connectivity between the SN and the putamen is disrupted in patients with RBD, when compared to controls (Ellmore et al., 2013). We thus next examined if the prodromal and control groups showed differing functional connectivity between left and right SN and the rest of the brain. This analysis did not reveal any significant clusters differentiating the prodromal and control groups.

## Discussion

The results demonstrate that functional connectivity between the different compartments of the BG is disrupted in individuals who are likely in the prodromal phases of PD, relative to age-matched controls. Subjects in the prodromal group showed reduced interhemispheric functional connectivity in each of the tested BG nuclei (putamen, caudate, pallidum) as well as reduced striato-thalamo-pallidal inter- and intrahemshpric connectivity. Connectivity changes were confined to striato-thalamo-pallidal structures in the BG and were not found between these structures and other cortical and subcortical regions.

Complex functional interactions between the different structures of the BG underlie its normal functioning, and disruption in this circuity, as demonstrated here in the prodromal group, is a central feature in mechanistic models of PD (Lanciego et al., 2012). The current results suggest that disruptions in striato-thalamo-pallidal functional interactions may occur already in the prodromal phase of PD, before the onset of the classical motor symptoms of the disease.

Our results join two other recent studies (Ellmore et al., 2013; Rolinski et al., 2016) in reporting intrinsic functional connectivity alterations which are manifested already in the prodromal phase of PD (Table 3). The current results differ from these earlier reports in our focus on connectivity within striato-thalamo-pallidal structures in the BG. The first study (Ellmore et al., 2013) utilized seed-based connectivity analysis, with the right and left SN as seed regions, and found differences between RBD patients and controls in functional connectivity between right SN and right parieto-occipital cortical regions and left SN and the left putamen (Ellmore et al., 2013). A second study used a data-driven approach and found reduced resting state-coactivation in RBD patients, relative to controls, in an extensive network encompassing the putamen, and prefrontal cortex. Thus, while these results both suggest that functional interactions of cortical and subcortical regions with the putamen are altered in prodromal PD, the methods used do not specifically focus on connectivity between the different input and output structures of the BG, as tested here. Our results specifically point to disruptions in local striato-thalamo-pallidal interactions, but not in functional connectivity between the BG and other cortical and subcortical regions, or between SN and striato-thalamo-pallidal structures. One challenge in comparing results between studies is that, as a progressive process, the prodromal phase of PD may be manifested differently in groups of subjects that are not matched closely in various attributes such as current clinical diagnosis and disease duration. It thus remains to be tested how functional connectivity within the BG and between the BG and other cortical and subcortical regions changes along the progression of the prodromal phase of PD.

**Table 3.**
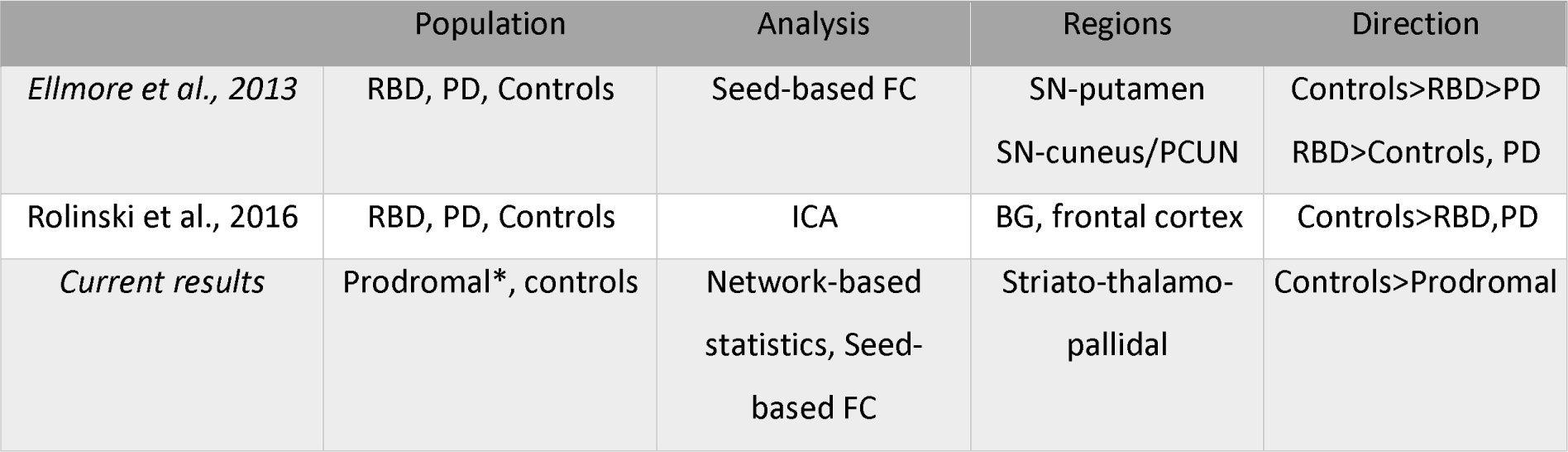
Comparison with previous results. The table compares the current results with those reported in two previous papers looking at resting-state fMRI alterations in patients with RBD. ICA, independent component analysis. All other abbreviations are as before. * composed of subjects with idiopathic RBD and hyposmia

Connectivity in the striato-thalamo-pallidal network did not significantly correlate with the prodromal group’s MOCA or UPDRS (part III, focusing on motor function) scores. Cognitive function is not considered a strong marker for prodromal PD, as it is not strongly predictive of phenoconversion from the prodromal to the clinical phase of the disease (Postuma et al., 2015b). Mild motor impairment, on the other hand, is moderately predictive of phenoconversion (Postuma et al., 2015b), however our current results did not reveal significant correlations between the motor parts of the UPRDS and functional connectivity in the striatothalamo-pallidal network. It should be noted, however, that the range of UPDRS part III scores displayed by the prodromal subjects analyzed here was limited (with more than 50% of subjects scoring either 0 or 1). Thus, at least in the current sample, the UPDRS may not have had the necessary sensitivity to detect mild motor impairment, as offered by other tests. Future work could focus more directly on links between imaging alterations in prodromal PD and markers of motor dysfunction.

While our results reveled consistent reductions in BG functional connectivity in the prodromal group relative to the control group, we did not find evidence pointing to alterations in the opposite direction (i.e., increases in functional connectivity in the prodromal group). Increased cortico-BG and thalamo-cortical connectivity has been reported in various populations of PD patients relative controls (Baudrexel et al., 2011; Gu et al., 2017; Kwak et al., 2010; Shen et al., 2017). It has been suggested that increases in functional connectivity may mirror excessive synchronization in cortico-BG circuitry (Baudrexel et al., 2011), as demonstrated with electrophysiological recordings (Brown, 2003). One possibility is that this form of aberrant connectivity only appears later in the course of PD. In fact, it was shown earlier that reductions in BG connectivity seen in RBD patients closely resemble those recorded from de novo PD patients (Rolinski et al., 2016). Thus, increased functional connectivity may constitute a pathophysiological feature appearing only later in the course of the disease.

An unresolved question is the degree to which markers of prodromal PD resemble (or possible continue) markers seen in asymptomatic risk populations. A recent study reported that asymptomatic LRRK2 mutation carriers, when compared with asymptomatic non-carriers show reduced functional connectivity between the caudal parts of the left striatum and the ipsilateral precuneus and superior parietal lobe, and increased connectivity between the right SN and bilateral occipital cortical regions. While these results suggest a more widespread pattern of alterations in cortico-BG functional connectivity relative to the ones reported here, a comparison between the studies should be done with caution, given the methodological differences between the studies and since the studied populations differed across multiple attributes. Future studies could look at the complete time-course of changes in cortico-BG functional connectivity from the risk and prodromal phases, through the early and then advanced stages of the disease. This will allow to uncover the dynamics of progressive changes in cortico-BG functional connectivity seen along the progression of the disease.

Several limitations in the current study should be taken into account. First, the prodromal PD cohort analyzed here consisted of patients with RBD or hyposmia. It is currently unclear if these markers of prodromal PD (Berg et al., 2015; Mahlknecht et al., 2015; Postuma and Berg, 2016) are associated with differing intrinsic functional interactions within the BG, and a larger sample would be needed to test this question. Second, the current results focus on a cross sectional single-time point comparison and it remains to be tested if the strength of BG connectivity could predict conversion from prodromal to clinical PD. Future adequately powered longitudinal studies will be needed in order to address this question (Heinzel et al., 2016).

Circuit- and network-based accounts of neurological (Sharp et al., 2014; Stam, 2014) and neuropsychiatric (Gunaydin and Kreitzer, 2016) disease states have been prevalent in recent years. Similar accounts have been proposed for PD, with data pointing to disruptions in multiple motor and non-motor brain systems and their interactions (Baudrexel et al., 2011; Dayan et al., 2012; Jaywant et al., 2016; Luo et al., 2014). With the current results in mind, it appears that a systems-level account for PD (Caligiore et al., 2016) may also offer new insight on its prodromal phase and the progression from the prodromal to the clinical phase of the disease.

## Acknowledgments

Data used in the preparation of this article were obtained from the Parkinson’s Progression Markers Initiative (PPMI) database (www.ppmi-info.org/data). For up-to-date information on the study, visit www.ppmi-info.org. PPMI – a public-private partnership – is funded by the Michael J. Fox Foundation for Parkinson’s Research and funding partners, including Abbvie, Avid Radiopharmaceuticals, Biogen, Bristol-Myers Squibb, Covance, GE healthcare, Genentech, GlaxoSmithKline, Lilly, Lundbeck, Merck, Meso Scale Discovery, Pfizer, Piramal, Roche, Servier, and UCB

## References

Baudrexel, S., Witte, T., Seifried, C., von Wegner, F., Beissner, F., Klein, J.C., Steinmetz, H., Deichmann, R., Roeper, J., Hilker, R., 2011. Resting state fMRI reveals increased subthalamic nucleus–motor cortex connectivity in Parkinson’s disease. NeuroImage 55, 1728–1738. doi:10.1016/j.neuroimage.2011.01.017

Berg, D., Postuma, R.B., Adler, C.H., Bloem, B.R., Chan, P., Dubois, B., Gasser, T., Goetz, C.G., Halliday, G., Joseph, L., Lang, A.E., Liepelt-Scarfone, I., Litvan, I., Marek, K., Obeso, J., Oertel, W., Olanow, C.W., Poewe, W., Stern, M., Deuschl, G., 2015. MDS research criteria for prodromal Parkinson’s disease. Mov. Disord. Off. J. Mov. Disord. Soc. 30, 1600–1611. doi:10.1002/mds.26431

Braak, H., Del Tredici, K., Rüb, U., de Vos, R.A.I., Jansen Steur, E.N.H., Braak, E., 2003. Staging of brain pathology related to sporadic Parkinson’s disease. Neurobiol. Aging 24, 197–211.

Brown, P., 2003. Oscillatory nature of human basal ganglia activity: Relationship to the pathophysiology of Parkinson’s disease. Mov. Disord. 18, 357–363. doi:10.1002/mds.10358

Caligiore, D., Helmich, R.C., Hallett, M., Moustafa, A.A., Timmermann, L., Toni, I., Baldassarre, G., 2016. Parkinson’s disease as a system-level disorder. Npj Park. Dis. 2, 16025. doi:10.1038/npjparkd.2016.25

Dayan, E., Inzelberg, R., Flash, T., 2012. Altered perceptual sensitivity to kinematic invariants in Parkinson’s disease. PloS One 7, e30369. doi:10.1371/journal.pone.0030369

Doty, R.L., Shaman, P., Kimmelman, C.P., Dann, M.S., 1984. University of Pennsylvania Smell Identification Test: a rapid quantitative olfactory function test for the clinic. The Laryngoscope 94, 176–178.

Eisensehr, I., Linke, R., Noachtar, S., Schwarz, J., Gildehaus, F.J., Tatsch, K., 2000. Reduced striatal dopamine transporters in idiopathic rapid eye movement sleep behaviour disorder. Comparison with Parkinson’s disease and controls. Brain J. Neurol. 123 ( Pt 6), 1155–1160.

Ellmore, T.M., Castriotta, R.J., Hendley, K.L., Aalbers, B.M., Furr-Stimming, E., Hood, A.J., Suescun, J., Beurlot, M.R., Hendley, R.T., Schiess, M.C., 2013. Altered Nigrostriatal and Nigrocortical Functional Connectivity in Rapid Eye Movement Sleep Behavior Disorder. SLEEP. doi:10.5665/sleep.3222

Gu, Q., Cao, H., Xuan, M., Luo, W., Guan, X., Xu, J., Huang, P., Zhang, M., Xu, X., 2017. Increased thalamic centrality and putamen-thalamic connectivity in patients with parkinsonian resting tremor. Brain Behav. 7, e00601. doi:10.1002/brb3.601

Gunaydin, L.A., Kreitzer, A.C., 2016. Cortico-Basal Ganglia Circuit Function in Psychiatric Disease. Annu. Rev. Physiol. 78, 327–350. doi:10.1146/annurev-physiol-021115-105355

Heinzel, S., Roeben, B., Ben-Shlomo, Y., Lerche, S., Alves, G., Barone, P., Behnke, S., Berendse, H.W., Bloem, B.R., Burn, D., Dodel, R., Grosset, D.G., Hu, M., Kasten, M., Krüger, R., Moccia, M., Mollenhauer, B., Oertel, W., Suenkel, U., Walter, U., Wirdefeldt, K., Liepelt-Scarfone, I., Maetzler, W., Berg, D., 2016. Prodromal Markers in Parkinson’s Disease: Limitations in Longitudinal Studies and Lessons Learned. Front. Aging Neurosci. 8. doi:10.3389/fnagi.2016.00147

Iranzo, A., Lomeña, F., Stockner, H., Valldeoriola, F., Vilaseca, I., Salamero, M., Molinuevo, J.L., Serradell, M., Duch, J., Pavía, J., Gallego, J., Seppi, K., Högl, B., Tolosa, E., Poewe, W., Santamaria, J., 2010. Decreased striatal dopamine transporter uptake and substantia nigra hyperechogenicity as risk markers of synucleinopathy in patients with idiopathic rapid-eye-movement sleep behaviour disorder: a prospective study. Lancet Neurol. 9, 1070–1077. doi:10.1016/S1474-4422(10)70216-7

Jaywant, A., Shiffrar, M., Roy, S., Cronin-Golomb, A., 2016. Impaired perception of biological motion in Parkinson’s disease. Neuropsychology 30, 720–730. doi:10.1037/neu0000276

Keuken, M.C., Bazin, P.-L., Backhouse, K., Beekhuizen, S., Himmer, L., Kandola, A., Lafeber, J.J., Prochazkova, L., Trutti, A., Sch?fer, A., Turner, R., Forstmann, B.U., 2017. Effects of aging on $$T_{1}$$ T 1, $$T_{2}^{*}$$ T 2⍰?, and QSM MRI values in the subcortex. Brain Struct. Funct. doi:10.1007/s00429-016-1352-4

Kwak, Y., Peltier, S., Bohnen, N.I., M?ller, M.L.T.M., Dayalu, P., Seidler, R.D., 2010. Altered Resting State Cortico-Striatal Connectivity in Mild to Moderate Stage Parkinson’s Disease. Front. Syst. Neurosci. 4. doi:10.3389/fnsys.2010.00143

Lanciego, J.L., Luquin, N., Obeso, J.A., 2012. Functional Neuroanatomy of the Basal Ganglia. Cold Spring Harb. Perspect. Med. 2. doi:10.1101/cshperspect.a009621

Luo, C., Song, W., Chen, Q., Zheng, Z., Chen, K., Cao, B., Yang, J., Li, J., Huang, X., Gong, Q., Shang, H.-F., 2014. Reduced functional connectivity in early-stage drug-naive Parkinson’s disease: a resting-state fMRI study. Neurobiol. Aging 35, 431–441. doi:10.1016/j.neurobiolaging.2013.08.018

Mahlknecht, P., Seppi, K., Poewe, W., 2015. The Concept of Prodromal Parkinson’s Disease. J. Park. Dis. 5, 681–697. doi:10.3233/JPD-150685

Marek, K., Jennings, D., Lasch, S., Siderowf, A., Tanner, C., Simuni, T., Coffey, C., Kieburtz, K., Flagg, E., Chowdhury, S., others, 2011. The parkinson progression marker initiative (PPMI). Prog. Neurobiol. 95, 629–635.

Nasreddine, Z.S., Phillips, N.A., Bédirian, V., Charbonneau, S., Whitehead, V., Collin, I., Cummings, J.L., Chertkow, H., 2005. The Montreal Cognitive Assessment, MoCA: a brief screening tool for mild cognitive impairment. J. Am. Geriatr. Soc. 53, 695–699. doi:10.1111/j.1532-5415.2005.53221.x

Postuma, R.B., Berg, D., 2016. Advances in markers of prodromal Parkinson disease. Nat. Rev. Neurol. 12, 622–634. doi:10.1038/nrneurol.2016.152

Postuma, R.B., Berg, D., Stern, M., Poewe, W., Olanow, C.W., Oertel, W., Obeso, J., Marek, K., Litvan, I., Lang, A.E., Halliday, G., Goetz, C.G., Gasser, T., Dubois, B., Chan, P., Bloem, B.R., Adler, C.H., Deuschl, G., 2015a. MDS clinical diagnostic criteria for Parkinson’s disease. Mov. Disord. 30, 1591–1601. doi:10.1002/mds.26424

Postuma, R.B., Gagnon, J.-F., Bertrand, J.-A., Marchand, D.G., Montplaisir, J.Y., 2015b. Parkinson risk in idiopathic REM sleep behavior disorder Preparing for neuroprotective trials. Neurology 84, 1104–1113. doi:10.1212/WNL.0000000000001364

Power, J.D., Barnes, K.A., Snyder, A.Z., Schlaggar, B.L., Petersen, S.E., 2012. Spurious but systematic correlations in functional connectivity MRI networks arise from subject motion. Neuroimage 59, 2142–2154. doi:10.1016/j.neuroimage.2011.10.018

Rolinski, M., Griffanti, L., Piccini, P., Roussakis, A.A., Szewczyk-Krolikowski, K., Menke, R.A., Quinnell, T., Zaiwalla, Z., Klein, J.C., Mackay, C.E., Hu, M.T.M., 2016. Basal ganglia dysfunction in idiopathic REM sleep behaviour disorder parallels that in early Parkinson’s disease. Brain aww124. doi:10.1093/brain/aww124

Rolinski, M., Griffanti, L., Szewczyk-Krolikowski, K., Menke, R.A.L., Wilcock, G.K., Filippini, N., Zamboni, G., Hu, M.T.M., Mackay, C.E., 2015. Aberrant functional connectivity within the basal ganglia of patients with Parkinson’s disease. NeuroImage Clin. 8, 126–132. doi:10.1016/j.nicl.2015.04.003

Sharp, D.J., Scott, G., Leech, R., 2014. Network dysfunction after traumatic brain injury. Nat. Rev. Neurol. 10, 156–166. doi:10.1038/nrneurol.2014.15

Shen, B., Gao, Y., Zhang, W., Lu, L., Zhu, J., Pan, Y., Lan, W., Xiao, C., Zhang, L., 2017. Resting State fMRI Reveals Increased Subthalamic Nucleus and Sensorimotor Cortex Connectivity in Patients with Parkinson’s Disease under Medication. Front. Aging Neurosci. 9. doi:10.3389/fnagi.2017.00074

Stam, C.J., 2014. Modern network science of neurological disorders. Nat. Rev. Neurosci. 15, 683–695. doi:10.1038/nrn3801

Whitfield-Gabrieli, S., Nieto-Castanon, A., 2012. Conn: a functional connectivity toolbox for correlated and anticorrelated brain networks. Brain Connect. 2, 125–141. doi:10.1089/brain.2012.0073

Zalesky, A., Fornito, A., Bullmore, E.T., 2010. Network-based statistic: identifying differences in brain networks. NeuroImage 53, 1197–1207. doi:10.1016/j.neuroimage.2010.06.041

Zeighami, Y., Ulla, M., Iturria-Medina, Y., Dadar, M., Zhang, Y., Larcher, K.M.-H., Fonov, V., Evans, A.C., Collins, D.L., Dagher, A., 2015. Network structure of brain atrophy in de novo Parkinson’s disease. eLife 4, e08440. doi:10.7554/eLife.08440

